# A novel dual-excitation pulse-amplitude-modulation fluorometer for investigating photosynthesis of plants

**DOI:** 10.1101/2024.04.18.589113

**Authors:** Martin Havelka, Ladislav Nedbal, Klaus Suhling, Jakub Nedbal

## Abstract

A modular instrument was developed to measure the fluorescence yield in plants subject to a combination of two harmonically-oscillating blue lights with independently controlled frequencies and phases.It uses the pulse-amplitude-modulation (PAM) method to measure the fluorescence yield independently of the plant irradiance. Compared to existing commercial instruments, it uses a higher measuring frequency (≈ 60 kHz) and higher measuring flash irradiance. This enables averaging over a number of subsequent measurement data points to achieve a higher signal-to-noise ratio. The manuscript describes the design, testing, and characterization of the operational limits of the instrument. It identifies its current weaknesses and makes recommendations for improvements. It is accompanied by supplementary materials containing the electronic schematics and the source code. The instrument was used to study a plant response to a mixture of two oscillating lights. It resulted in an excellent signal-to-noise ratio of the measured fluorescent yield. The measurements clearly demonstrated that the fluorescence yield of a plant subject to a combination of two harmonically-oscillating lights is not the same as the sum of the responses to the two oscillating lights applied independently. The observed non-linearity leads to the important conclusion that the time- and frequency-domain cannot be connected by a Fourier transform. Therefore, the frequency-domain approach will yield novel information that is not redundant to the well-established time-domain measurements.

## I. INTRODUCTION

Photosynthesis of plants and algae provides free energy for all major forms of life including humans, sustains the oxygenic atmosphere, and stabilizes the climate by capturing CO_2_ ^1^. Under optimal conditions, most of the sunlight absorbed by chlorophyll drives this unique process. Yet, a variable part of the captured energy is also emitted as chlorophyll fluorescence or dissipated as heat. Measuring chlorophyll fluorescence offers a non-invasive technique, which provides insight into dissipative pathways competing with photosynthesis, its regulatory mechanisms, and indirectly the health or stress state of the organism^2^. It is notable that chlorophyll fluorescence response is highly complex and non-linear compared to fluorescence emission from most other inanimate or biological chromophores like fluorescent dyes or proteins. Equally, it is often more informative.

The unifying basis for measuring photosynthetic parameters by fluorescence is through its ‘quenching’, whereby the photochemistry and the heat dissipation compete with fluorescence for the available harvested light energy.^3^ The high efficiency of photosynthesis in healthy plants exposed to sub-saturating light levels is reflected by a chlorophyll fluorescence yield that is reduced to minimal levels by the ‘photochemical quenching’. In contrast, when the efficiency of photochemical reactions is reduced by high light levels that saturate the downstream electron transport pathways, the ‘photochemical quenching’ is ineffective and chlorophyll fluorescence yield reaches its maximal levels.

At a high light irradiance, the imbalance between the excessive photochemical energy conversion in the reaction centers and the limited capacity of the coupled oxidation-reduction reactions can produce harmful radical oxygen species. This and other forms of photoinhibition in high light can be reduced by a number of protective mechanisms that fall under the umbrella term ‘nonphotochemical quenching’ (NPQ)^4^. The surplus absorbed light energy is dissipated as heat or reduced by decreasing the number of chlorophyll molecules that supply the reaction centers with excitation energy, both reducing not only the photochemistry but also chlorophyll fluorescence. The interplay between photosynthetic organism surface irradiance, the photochemical and nonphotochemical quenching, and other mechanisms thus leads to the highly non-linear response observed in the chlorophyll fluorescence of photosynthetic organisms.

Therefore, a number of measurement principles and instruments emerged, that use fluorescence sensing to the benefit of photosynthesis research and its applications^5^. The methods range from sensing sun-induced chlorophyll fluorescence from space or airborne platforms^6^ to measuring the emission from individual cells or chloroplasts^7^. Most of the proximal methods use a dim narrow-band measuring light that excites fluorescence without changing the photochemical status of the photosynthetic organism. This is mostly achieved by exciting fluorescence with weak microsecond-short measuring flashes and synchronous detection. The synchronous detection eliminates interference of fluorescence that is excited by slowly changing harmonic light. This so-called actinic light generates substantial photochemical activity in plants, unlike the dim measuring flashes. It is applied to probe the dynamics of the photochemical and nonphotochemical quenching. A combination of dim, actively modulated measuring light pulses and strong actinic light is also used in the widely applied pulse amplitude modulation (PAM) technique that was introduced to measure the quantum yield of photochemical conversion in photosynthesis and other photosynthetic parameters^3^. The PAM method has been commercially exploited by various suppliers of field and laboratory instruments.

The PAM instruments were primarily designed to capture fluorescence transients appearing in plants in response to dark-to-light transitions and/or light pulses. Nedbal and Březina proposed to study the fluorescence dynamics in the frequency domain^8^ and, since then several commercial instruments have been developed. They approximate harmonic modulation of the actinic light, which drives the photochemical reactions in photosynthesis, by step changes. Such instruments were employed in several pilot studies^9–11^ that not only brought new insights into photosynthetic dynamic and regulation but also identified multiple deficiencies of these PAM instruments when applied in the frequency-domain. Namely, the frequency-domain studies require accurate sine-modulated light that is only poorly approximated by multiple-step light changes generated by the instruments that were originally designed to measure fluorescence transient responses to square-light modulation. This limitation is particularly severe, e.g., when the light steps are large relative to the low amplitude of the light modulation. Fourier analysis and Bode plots, showing the phase and magnitude dependence on the modulation frequency of the signals acquired by these instruments, are then strongly affected by this artifact. Another inherent property of the existing instruments is the requirement to measure chlorophyll fluorescence yield in dark-acclimated plants. To achieve this, the measuring light flashes must be relatively weak and, consequently, the signal-to-noise ratio can be low which may limit measuring plants from a distance, e.g., in relevant greenhouse or field applications. This is not the case when the photosynthetic dynamic is probed in the light-acclimated plants exposed to the harmonically-modulated actinic light of variable frequency. Then, the measuring flashes can be much stronger and applied more frequently for improved signal-to-noise ratio. Elimination of the dark adaptation from the experimental protocols is another important advantage of studying photosynthetic dynamics in the frequency domain. Further, the signal processing of the commercial PAM instruments cannot filter out the high-frequency noise components without deforming the signal’s fastest components that may appear in response to the step-wise light changes. In contrast, the pilot studies^10,11^ showed that the plant responses to smooth, harmonically modulated light consist of the fundamental harmonics of the frequency that is the same as light modulation and only a small number of upper harmonics. This information can be used for effective signal filtering that can further increase the signal-to-noise ratio of instruments that would be designed specifically for frequency-domain measurements.

This paper presents a new instrument, further called Harmonizer, that may serve as a base for easy and inexpensive reproduction and further development of PAM-type instruments for the frequency-domain characterization of plants. The Harmonizer is described here in detail, its operating limits are characterized, and further steps toward increasing the application range of the Harmonizer are suggested. The performance of the Harmonizer is demonstrated here by probing the linearity of the plant responses to the harmonically modulated light. Towards this goal, the instrument generates simultaneously two harmonic light components that can be modulated with different periods and amplitudes. This will, in future studies, probe the range of periods and amplitudes that prompt an additive linear response of the investigated plant. This is essential for identifying the range in which the photosynthesis response to complex fluctuating light environments in plant canopies can be predicted from the responses to the constitutive harmonic components^11^.

## II. INSTRUMENT DESCRIPTION

The instrument is made of a microprocessor development board (Raspberry Pi, RPi) with an attached touch screen, a microcontroller development board (Arduino), and five custom-build electronics circuit boards (Fig. 1). The RPi served as the main controller of the Harmonizer, facilitated the interaction with the user and stored the measurement data. It was running a custom Python script, which initiated the electronic elements of the Harmonizer and started the measurement. The initiation involved firstly setting up the output voltages on the digital-to-analog converters (DAC) and the phases and frequencies on the direct digital synthesis (DDS) circuits. The DACs adjusted the oscillation amplitude and the constant offset of the harmonically modulated lights. A separate DAC set the amplitude of the measuring light flashes. The instructions to the DAC and DDS circuits were sent through a Serial Peripheral Interface (SPI) bus. The second part of the initiation involved setting the experiment duration and DDS clock divider ratio in the Arduino through an Inter-Integrated Circuit (I^2^C) bus. At this point, the Arduino microcontroller took over and ran the measurement protocol. It sent a clock signal to the DDS, reset the DDS to start harmonic oscillations with the programmed initial phase, started sending regular measuring light flashes, and synchronous conversion signals into the analog-to-digital converter (ADC). The SPI output of the ADC was connected to a demultiplexer (DEMUX), which was also controlled by the Arduino. It sent the ADC output data into one of the two present static random access memories (RAM). The RAMs were connected to a multiplexer (MUX), which alternated their connection to a common SPI port on the RPi. The MUX ensured the RAM, which was not being written to, was available to the RPi for readout. Once the Arduino filled up the other RAM with measurement data, it flipped the MUX/DEMUX circuit and send a pulse (full) to the RPi. The (full) signal was an interrupt instructing the RPi to read the content of the available RAM and concatenate it with the previously read RAM content. This cycle was repeated until the end of the measurement. At this point, the RPi took back control of the circuit and shut down the lights. Meanwhile, the Arduino became idle until the next measurement instruction. In the following paragraphs the modules are described in more detail.

**FIG. 1.**
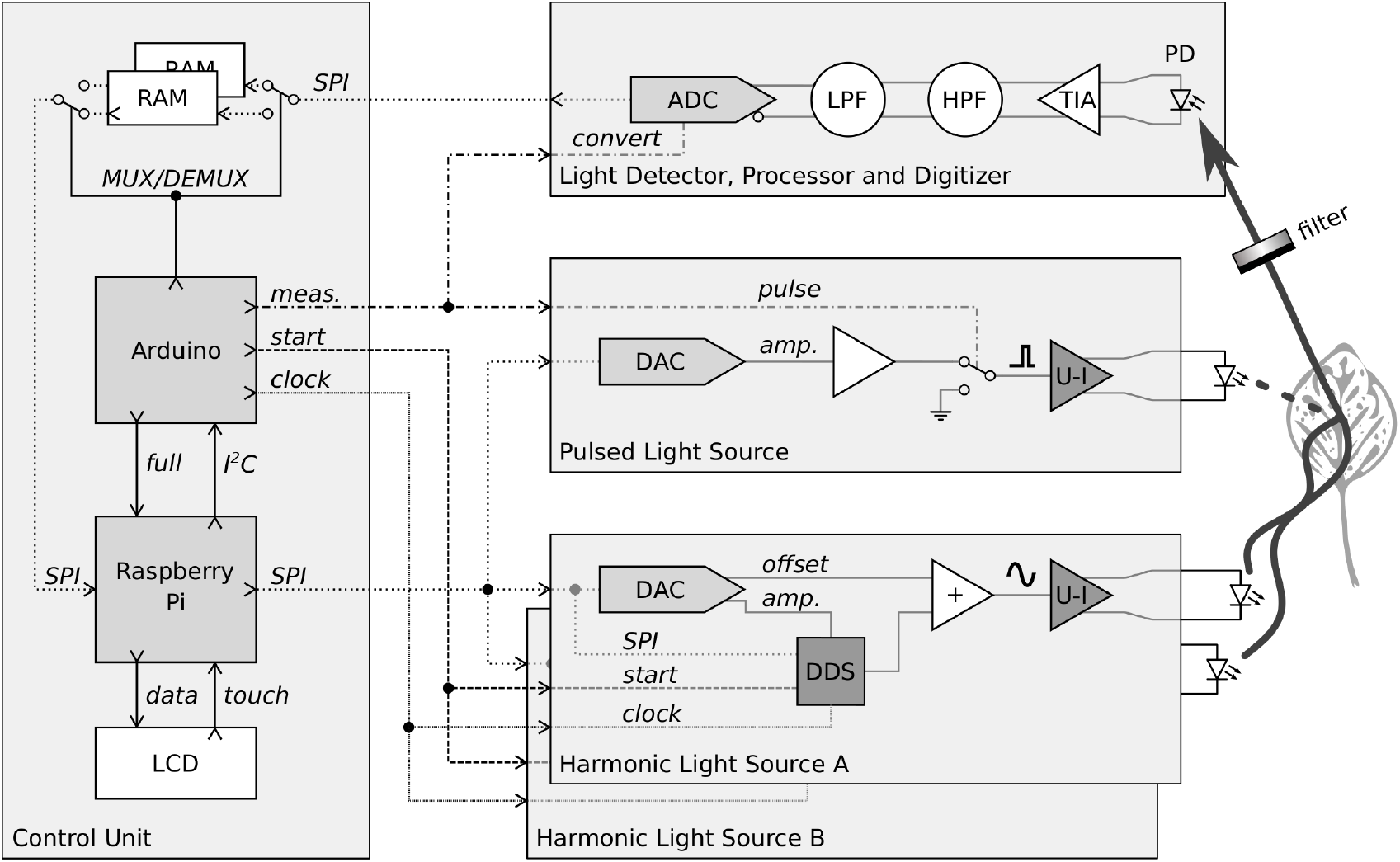
Schematic diagram of the Harmonizer. A Raspberry Pi microprocessor board controlled the operation of the Harmonizer and stored the measurement data. An Arduino microcontroller controlled the uninterrupted real-time data transfer into buffer memories (RAM) during the measurement. Two harmonic light sources produced the actinic light. The digital-to-analog converter (DAC) controlled the offset and the amplitude of the light. Direct digital synthesis (DDS) controlled its frequency and phase, summing amplifiers (+) added the offset. A voltage to current amplifier (U-I) drove the LED. A pulsed light source produced the measuring flashes with a DAC-programmable amplitude. The chlorophyll fluorescence of the leaf was spectrally filtered and detected by a photodiode (PD) with its current amplified by a transimpedance amplifier (TIA). The fluorescence resulting from the measuring flashes was separated from the fluorescence excited by the harmonic lights by a series of high-pass filters (HPF) and low pass filters (LPF) before being digitized by an analog-to-digital converter (ADC). The data stream was stored into one memory (RAM) while the other memory was available for reading into the Raspbery Pi

### A. Microprocessor

The microprocessor board was Raspberry Pi 3 Model A+ (SC0130, The Pi Hut, Haverhill, UK) with 1.4 GHz ARM Cortex-A53 processor and 512 MB RAM and a 16 GB secure digital (SD) card (The Pi Hut) for the operating system and storage. The RPi had NOOBS Raspberry Pi operating system with graphical interface installed, which was kept regularly updated. The human interface was facilitated by remote access from a personal computer via secure shell (SSH) or directly through the connected 5” touchscreen (WAV-18396, The Pi Hut). The RPi had Python 3.9 installed to run the Harmonizer script main.py.

### B. Microcontroller

The microcontroller board was Arduino UNO (A000066, Farnell, Leeds, UK) with a 16 MHz ATmega328P microcontroller. The microcontroller code was programmed with all timer interrupts disabled and all timings determined by the code structure and delays introduced by no operation (NOP) instructions. The code often used direct register manipulation to minimize delays. The Arduino code served two roles, one was to accept initiation instructions via I^2^C bus from the RPi and the second was to execute the measurement protocol^12^. The initiation process involved setting the frequency of the master clock driving the circuits synthesizing the harmonic waveforms and setting the duration of the measurement in seconds. Once the setup was complete, the Arduino started the measurement. The measurement involved three levels of timings. At the fastest rate of 59.88 kHz, the Arduino sent pulses to drive the measuring light flashes, to control the ADC conversion, and the data transfer. At 1 Hz frequency, the Arduino switched the multiplexer/demultiplexer pair to swap the RAMs for storing the measurement data. It initialized the new RAM for writing starting from the lowest address. It also sent an interrupt pulse to the RPi to inform it of the new data being available for readout. The code ensured there was no delay introduced into the measuring pulse train during this transition. Finally, the Arduino counted the number of elapsed seconds to end the measurement at the right time. At the end, it set all outputs to the default settings and it stopped any further activity.

### C. Control Unit

The control unit connected all the modules together. In addition, it contained the power supply units and the RAMs with the multiplexer/demultiplexer pair. All modules, except the light detector module, were connected to the control unit board through stackable pin headers and faced outwards. The outward arrangement made the Harmonizer larger than necessary, but easier to debug. The power supplies consisted of three DC-DC converters (ISE0505A) producing separate analog voltages and a ground (+1.8 V, +3.3 V, +4.5 V, 4.5 V) and a digital supply voltage and a ground (+4.5 V). The source of the current to the power supplies was the RPi. The multiplexer/demultiplexer circuit was made of four three-way three-state switches (74LVX244), which switched the connections of the two RAMs’ (23LC1024) SPI buses to either the RPi, Arduino, or the analog-to-digital converter (ADC). The connection to the RPi facilitated the reading out of the RAM content, the connection to Arduino enabled writing to the RAM by providing the clock and chip select signals, but also data to initialize the burst writing sequence. Once initialized, the data input was connected to the ADC to fill up the memory with the measurement data.

### D. Harmonic Light Source

The harmonic light source consisted of two parts, a lownoise mixed signal circuit, which generated the driving voltage, and a high-current LED driver. The module contained two programmable parts, a two-output digital-to-analog converter (DAC) and a direct digital synthesis (DDS) chip connected to the RPi through an SPI bus. The SPI bus chip select signals for the DAC and DDS were connected to two output pins of the RPi through a three way solder bridge. Depending on the connection in the solder bridge, two otherwise identical modules connected in parallel could be addressed separately by the RPi. The DAC (MCP48CVB02) set the constant offset to the LED current through one output and the oscillation amplitude through its second output. The amplitude signal was fed to the DDS chip (AD9834), which used it as a reference current sink. The DAC output voltages and the DDS phase and frequency were programmed prior to the measurement start by the RPi. The DDS also required a clock input, which was provided by the Arduino at a frequency programmed by the RPi. The Arduino also reset the DDS to the programmed starting phase right before the start of each measurement. This ensured consistent performance, but also synchronized the phases in the two otherwise independent light source modules. The output of the DDS and the offset voltage of the DAC were summed in a summing operational amplifier (MCP6286) and sent onto a voltage-to-current converter (U-I) with an LED connected to its output. The U-I circuit used an operational amplifier (OPA728) and a power bipolar transistor (2SC6097) to supply current to the LED in a linear proportion to the controlling input voltage. Each module contained two U-I converters, to drive up to two diodes, but only one was assembled and used. The LEDs were connected through short lengths of coaxial cables to the SMA connectors on the module boards. The modules also contained a position to solder a DC power input connector, which was there to supply current to the LEDs. This connector was unused, too. Instead the power was shared by the pulsed light source module through the pin header connectors connecting all modules together.

### E. Pulsed Light Source

The pulsed light source module used a single output of the same DAC (MCP48CVB02) described above. This output set the amplitude of the measuring light flashes. The output voltage was fed through an analog switch (74LVC1G3157) controlled by the Arduino. The output of that switch was fed to the same U-I circuit described above. The DC power supply connector was assembled on this board. It was used to supply current to all three LED driver modules from a 5 V 1 A plug-in adapter.

### F. Light Detector, Processor, and Digitizer

The light detector, processor, and digitizer module was the most complex part of the Harmonizer. It contained a photodiode amplifier and a band pass filter. The circuit was designed to provide a voltage output proportional to the fluorescence arising from the measuring light flashes only. The fluorescence intensity arising from the lower frequency harmonically modulated actinic light was filtered out by this circuit. The module also contained an analog-to-digital converter (ADC) to digitize the measured fluorescence pulse amplitude and store it into the RAM on a command from the Arduino. The module circuitry was differential to minimize any noise interference. It consisted of a transimpedance amplifier (TIA) (ADA4817-1) for the photodiode (BPW34). This was followed by an active amplifying high-pass filter (HPF) (AD8066) and a passive high-pass filter. The RC filter components were empirically tweaked to provide optimal performance on inspection with an oscilloscope. The role of the high-pass filters was to suppress electrical signals arising from the fluorescence excited by the harmonically oscillating light, yet pass through the signal arising from the short measuring flashes. The high-pass filters were followed an amplifying active anti-aliasing low-pass filter (LPF) (THS4551) and a passive low-pass filter. The RC components of the active filter were also tweaked to optimize its performance. The output of the passive filter was fed into an ADC (MCP33131D-05) with an SPI output. The conversion signal and the SPI bus control were provided by the Arduino to sample the voltage at the peak of the measuring flash fluorescence response. This module was the only one connected to the control unit board through a short (15 cm) flexible ribbon cable instead of stackable connectors. This allowed attaching it to the optomechanical leaf clip assembly positioned on the top of the Harmonizer.

### G. Optomechanical Assembly

The Harmonizer electronics was sandwiched between two 6 mm-thick plywood boards and held together by M4 nylon stand-offs (Fig. 2**A**). The top plywood board had two openings for flat cables, one to connect the LCD touchscreen and the other to connect the light detector module. The touchscreen was fixed directly to the plywood by screws. The light detector module was screwed to a 3D printed holder with a leaf clip and fixed to the plywood by three stand-offs (Fig. 2**B**). The purpose of the holder was to position the leaf surface, the three illuminating LEDs, their collimator lenses, an optical filter, and the light detector module in a spatial arrangement ensuring efficient and homogeneous sample illumination and fluorescence light detection. The 3D printed holder was made of black polylactic acid (PLA) (3D People, London, UK). The leaf clip was made of a separate 3D-printed part connected to the main assembly by a spring-loaded steel hinge. The LEDs came mounted on an aluminium-backed boards (Cree XP-E Royal Blue). 25 cm-long coaxial cables with male SMA connectors on one end were soldered to the LEDs. The LEDs were then screwed to 36 mm-diameter 10 mm-thick aluminium heatsinks with heatsink compound at the interface. Narrow beam collimator lenses 120/223 by Polymer Optics (181-0728, RS Components, Corby, UK) were fixed over each LEDs by a thin double-sided sticky tape. The assemblies consisting of the LED, heatsink, supply cable, and lens were screwed to the 3D printed assembly in the prepared locations to provide three illumination sources. Two LEDs provided the harmonic actinic light and one delivered the measuring light flashes onto the leaf surface. A 20-mm diameter 5° narrow beam collimator lens was fixed to the printed circuit board in front of the photodiode on the light detector module to greatly increase the amount of collected fluorescence light. An interference band-pass filter with 670 nm center wavelength and 40 nm full-width at half-maximum (670DIR25, Knight Optical, Harrietsham, UK) was glued into the 3D printed holder opening using a dental casting adhesive (Twinsil, Picodent, Wipperfürth, Germany). This filter excluded the excitation light from reaching the photodiode and transmitted the fluorescence mainly produced by the photosystem II reaction centers. The light detector module was attached to the 3D-printed holder by three stand-offs. These stand-offs were fixed by screws to the top plywood board. The three LED assemblies were connected through their SMA connectors to the respective LED drivers. The remaining modules were connected together by header pin stacking connectors and by stand-offs to the plywood bases to create a stable structure. The dimension of the plywood bases of the Harmonizer was 18 ×18 cm^2^. The footprint was enlarged by 10 cm in width due to the extending LED supply cables and by 1 cm in depth due to one protruding LED heatsink. The overall height of the Harmonizer was 16 cm.

**FIG. 2.**
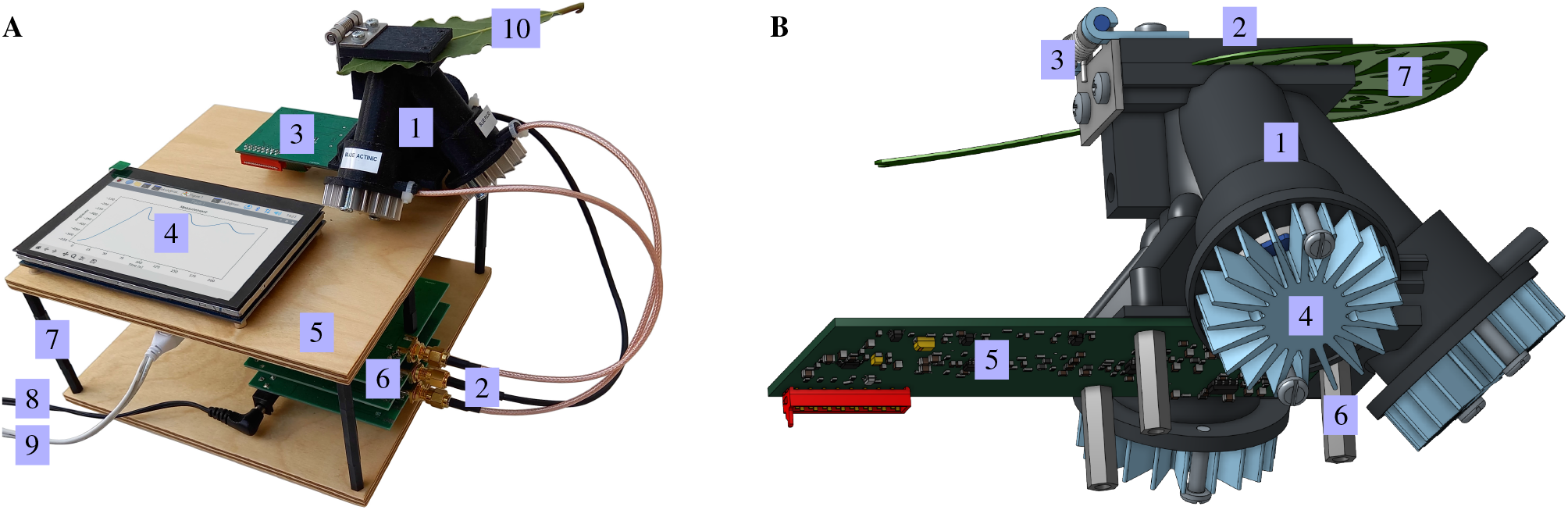
Assembled Harmonizer. **A** Photograph of the operating Harmonizer. (1) Optomechanical assembly with the leaf clip is where the measurements were taking place. (2) Coaxial cables connected the two harmonic light sources and one pulsed light source modules to the LEDs. (3) Light detector module PCB was fixed to the optomechanical assembly. (4) The touchscreen LCD display shows the measurement. (5) The top plywood board was connected to the bottom one in the corners by 74 mm-high standoffs. The power supply was supplied (8) to the pulsed light source module and (9) to the RPi. (10) The leaf was held inside the optomechanical assembly by the leaf clip. **B** Rendering of the assembled (1) 3D-printed optomechanical holder. The 3D-printed (2) leaf clip was connected to the holder through a (3) spring-loaded stainless steel hinge. Three LED assemblies with lenses and (4) star-shaped aluminium heatsinks were attached to the holder by two screws each. (5) The printed circuit board of the light detection module was fastened by three (6) M3 18 mm-long standoffs, also used to mount the assembly to the top plywood surface of the Harmonizer. (7) Leaf was clipped between the clip and the holder for the duration of the measurements.

### H. Harmonizer Application to Plant Fluorescence

The Harmonizer measured the fluorescence response on a 10 mm diameter circular area of leaf surface held in place by a spring-loaded leaf clip. The operation was controlled by a Python script, which executed the measurement and is described in Appendix A. Briefly, the script could run with any number of inputs, which set the experimental parameters (amplitude, phase, and frequency of harmonic light, amplitude of measuring light flashes, experimental duration, experiment title, and whether to produce an output graph). The script executed the experiments based on the inputs and created a binary file with the measurement data and the optional graph. The output filenames contained all the experimental parameters (metadata) for record.

## III. HARMONIZER CHARACTERISATION

The Harmonizer performance was characterized by a set of calibration measurements intended to establish its operating limits. An oscilloscope (SDS1202X-E, Siglent, Wokingham, UK) was used to record waveforms of electronic signals and a power meter (PM100A, Thorlabs, Ely, UK) with a calibrated power diode (S120VC, Thorlabs) were used to calibrate the sample irradiance.

### A. Measuring Flash Characterization

The PAM measurement principle employed in the Harmonizer measured the fluorescence excited by the measuring light flashes while excluding the background fluorescence excited by the harmonically modulated actinic light sources. The measuring light flashes intensity and frequency were constant and any changes to the detected fluorescence signal were the result of the change of the fluorescence yield of the sample. The harmonic light drove the changes in the sample but on its own did not contribute to the measured fluorescence directly. The Harmonizer used electronic frequency filters to separate the measuring flash-induced fluorescence from the harmonically modulated light-induced fluorescence. This section presents a set of experiments that verified the correct functionality of the Harmonizer in PAM measurements and determined its operating range.

The first experiment characterized the electronic signals associated with the measuring flashes and the detection of the resulting fluorescence (Fig. 3). A green fluorescent slide (Chroma, Bellow Falls, VT, USA) was used for these calibration measurements instead of a leaf as it had a linear response of fluorescence to excitation light intensity and exhibited high photostability. The oscilloscope was used to measure the current through the LED, the photodiode pre-amplifier output voltage and the differential voltage at the input to the analog-to-digital converter. The measurements show the shape of the signals and their relations in time and amplitude (Fig. 3) and that the light flash duration was just under 2 μs. The photodiode output and ADC input signals were delayed by a few hundred nanoseconds in respect to each other and the LED current pulse. The signal on the ADC input suffered from significant ringing, which however largely decayed by the time of the next pulse. The ADC was sampling its input for ≈ 3 μs (Fig. 3**B**). This sampling time was optimized to maximize the ADC reading. The characterization measurements revealed opportunities for improvements to the electronics signal processing circuits to decrease the ringing and to expand the pulse duration to match the ADC sampling window. Implementing these changes could improve the sensitivity and the linear operating range of the instrument.

**FIG. 3.**
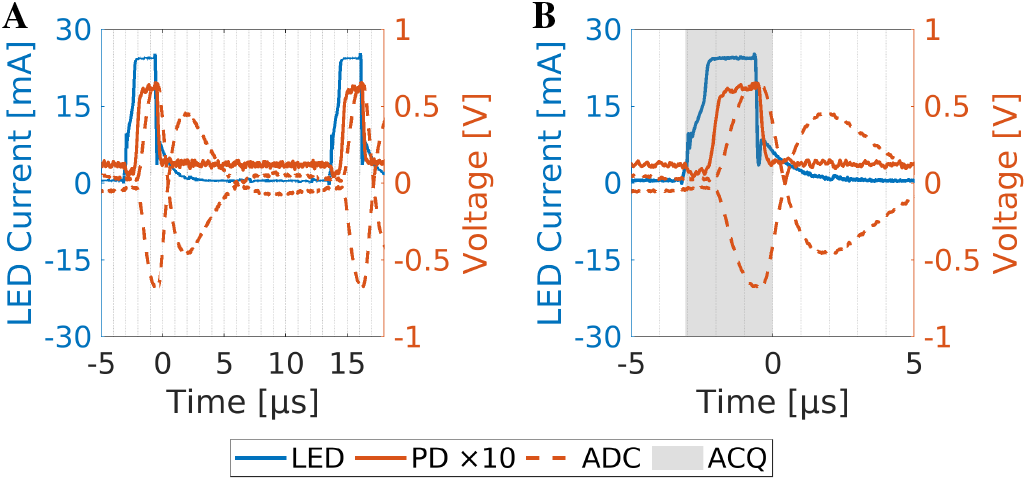
Electrical characteristics of measuring flashes. Oscilloscope traces show the current through the LED (LED, blue line), photodiode pre-amplifier output voltage, and the input into the analog-to-digital converter. The photodiode pre-amplifier output voltage (PD ×10, orange solid line) was amplified ten times in the graph for clarity. The differential ADC input was drawn by an orange dashed line (ADC). Grey box highlighted the acquisition time range (ACQ), where the ADC was sampling the input signal. **A** shows two pulses of the same measurement to demonstrate that signals were relaxed before the subsequent pulse appeared. **B** Shows a detail of a single pulse with the ADC sampling time windows in gray.

Next, the linear range of the Harmonizer was established in a set of experiments measuring its response to gradually increasing measuring light flash settings (Fig. 4). Three parameters - LED current, photodiode pre-amplifier output, and Harmonizer measured output - were examined during these measurements to establish where non-linearities arose in the system. The LED current and photodiode pre-amplifier output voltage were sensed using the oscilloscope and the measured output was the measurement made by the Harmonizer ADC. The fluorescent slide was used as the calibration sample and a semitransparent tape was stuck to its surface when needed to attenuate the passing light below the saturation point of the light detector circuit. The data was acquired with measuring flash settings between 7 and 100. A linear regression was calculated for the measurement data to determine the limits for the linear dependency on the measuring flash setting. The results show: (1) The LED current was linear in the entire tested range (Fig. 4**A, B**). (2) The photodiode pre-amplifier output voltage was linear between the settings of 7 to 40. At higher settings, the voltage started to fall below the linear regression line. This was probably due to the heating of the LED chip, which decreased its light-emission efficiency. At settings of 6 and below, there was no measurable signal on the photodiode pre-amplifier output. At settings of 50 or above the the output signal slowly diverted from the linear dependence. This deviation did not mean that any measurements with flashes at higher settings than 40 were invalid or nonlinear. It meant that it was no longer possible to assume that doubling the setting would lead to doubling of the measuring flash light energy. (3) The linearity of the signal processing and analog-to-digital converter circuitry was ascertained by comparing the Harmonizer measured output to the measuring light flash settings (Fig. 4**C, D**). Two samples with high and low fluorescence intensities were used, differing by light-attenuating tape layer attached to the fluorescent slide surface. This experiment showed a more limiting non-linearity and signal degradation at higher fluorescence intensities compared to (2). One observation was that once the Harmonizer measurement exceeded ≈ 800 it became highly non-linear and non-monotonous. This limited the safe maximum fluorescence range to ≈ 600 to stay well within the monotonous range and avoid a large measurement error. Lower fluorescence signals were reasonably linear and became strictly linear below ≈ 100.

**FIG. 4.**
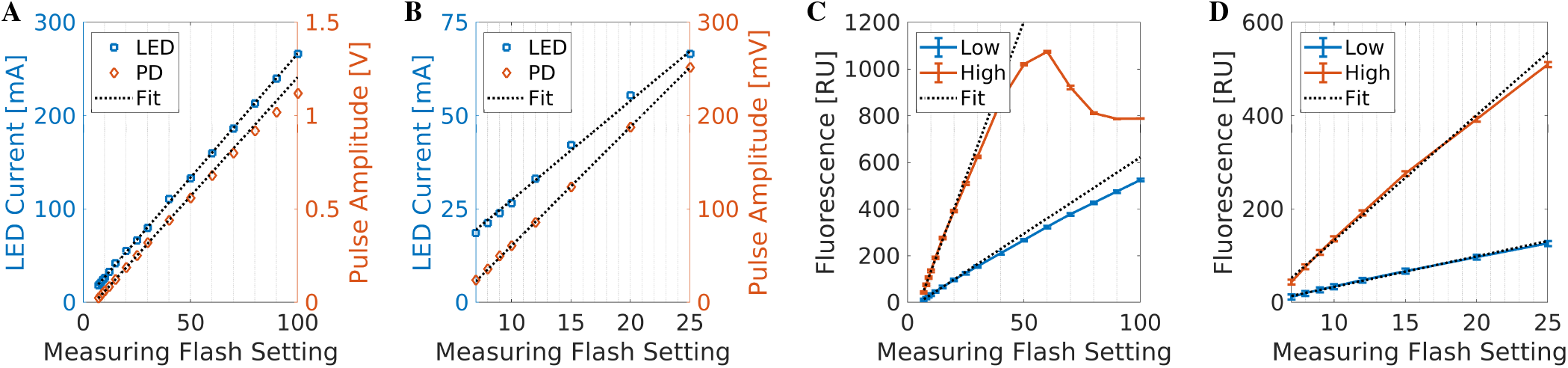
Measuring flash linearity range. **A, B** Flash amplitude linearity verification, where LED current amplitude (LED, blue square) and photodiode pre-amplifier voltage amplitude (PD, orange diamond) are plotted against the measuring flash setting. The linear fit (black dotted line) is the regression through all data points of the LED current and through the first eight points of the photodiode pre-amplifier voltage, where the response is linear. **A** Measuring flash setting was varied between 7 and 100. Settings of 50 and above show deviation of the photodiode pre-amplifier voltage below the regression line. This suggests that the LED light output becomes non-linear with supply current at settings higher than 50. **B** Same as **A**, with the exception of measuring flash setting range being between 7 and 25, where both the LED current and photodiode pre-amplifier outputs were linear with respect to the measuring flash setting. **C, D** Fluorescence measurements as a function of the measuring flash settings. Samples of low (blue line) and high (red line) fluorescence intensities were measured. The results show that signals in excess of 600 became unrepresentative due to amplifier saturation and resulting signal degradation. The fluorescence measurement was linear for values below ∼ 100, but deviated from the linear dependence between 100 and 600. Error bars represent the standard deviations of the each data point.

The linearity measurements found that the measurement flash intensity was reasonably linear up to the maximum examined setting value of 100 and were completely linear at settings of 40 and below. Any setting below 7 meant the flashes were turned off. Harmonizer measurements with values below 100 were strictly linear in respect to the fluorescence yield of the sample. Signals above 100 have shown a slightly non-linear relation to the sample fluorescence and signals above 600 were found to be unreliable and should always be avoided.

### B. Harmonic Light Effect on the Measurement

The Harmonizer was designed to suppress fluorescence arising from the harmonic light sources and only detect the fluorescence arising from the measuring flashes. The experiments described below were designed to find the operational range where it met the above criteria. The first experiment was designed to answer the question if and under what conditions the harmonic light-excited fluorescence signal reached the analog-to-digital converter and thus perturbed the measured results. Since the measuring flash fluorescence signal was separated from the harmonic light-excited fluorescence signal by the means of high-pass electronic filters, it was clear that any bleed-through of harmonic light-excited fluorescence was going to happen mainly at higher frequencies, where the filter suppression was going to be less effective. The experiment was done with constant harmonic light amplitude and measuring flash amplitude settings. The frequency of the harmonic light was varied between 1 kHz and 10 kHz. The sample was either a fluorescent slide, as above, or a leaf (*Laurus nobilis*). An experiment using the fluorescent slide and no harmonic light was performed as a control. The results are summarized in Fig. 5. In the fluorescence slide measurement (Fig. 5**B**), no modulated signal was apparent in the data for any of the frequencies. However, their Fourier transforms (Fig. 5**A**) showed signs of harmonic light-induced fluorescence, which appeared as peaks in the frequency space at the modulation frequencies and their higher harmonics. These peaks were small for the harmonic light with 10 kHz frequency, they dropped just above the noise floor at 2 kHz, and they became lost in the noise floor at 1 kHz. With the leaf (Fig. 5**C**), the amplitudes of the Fourier spectrum peaks at the modulation frequencies were higher at the 1 kHz and 2 kHz frequencies of the harmonically modulated light compared to the fluorescent slide sample in (Fig. 5**A**). The peak amplitudes at the modulation frequencies were higher than their harmonics peaks, unlike in the controls, where they were similar. This increase could be explained by an actinic light effect on the fluorescence yield of the leaf at those frequencies. Also apparent was the fluorescence induction of the leaf during the experiment (Fig. 5**D**), which was absent in the results from the fluorescent slide sample (Fig. 5**B**). In the last control measurement, the fluorescent slide was used without any harmonic light illumination and showed flat responses in the measured fluorescence (Fig. 5**F**) and its Fourier transform (Fig. 5**E**). In summary, this experiment concluded that the Harmonizer completely suppressed all fluorescence excited by the harmonically modulated lights at frequencies of 1 kHz and lower. This meant that any detected fluorescence change at these lower frequencies was entirely caused by the change to the fluorescence yield of the sample. At the same time, there was a small contribution of cross-talk of the harmonic light into the measurement at frequencies of 2 kHz and above. This experiment established the maximum limit for the Harmonizer harmonic light frequency to 1 kHz.

**FIG. 5.**
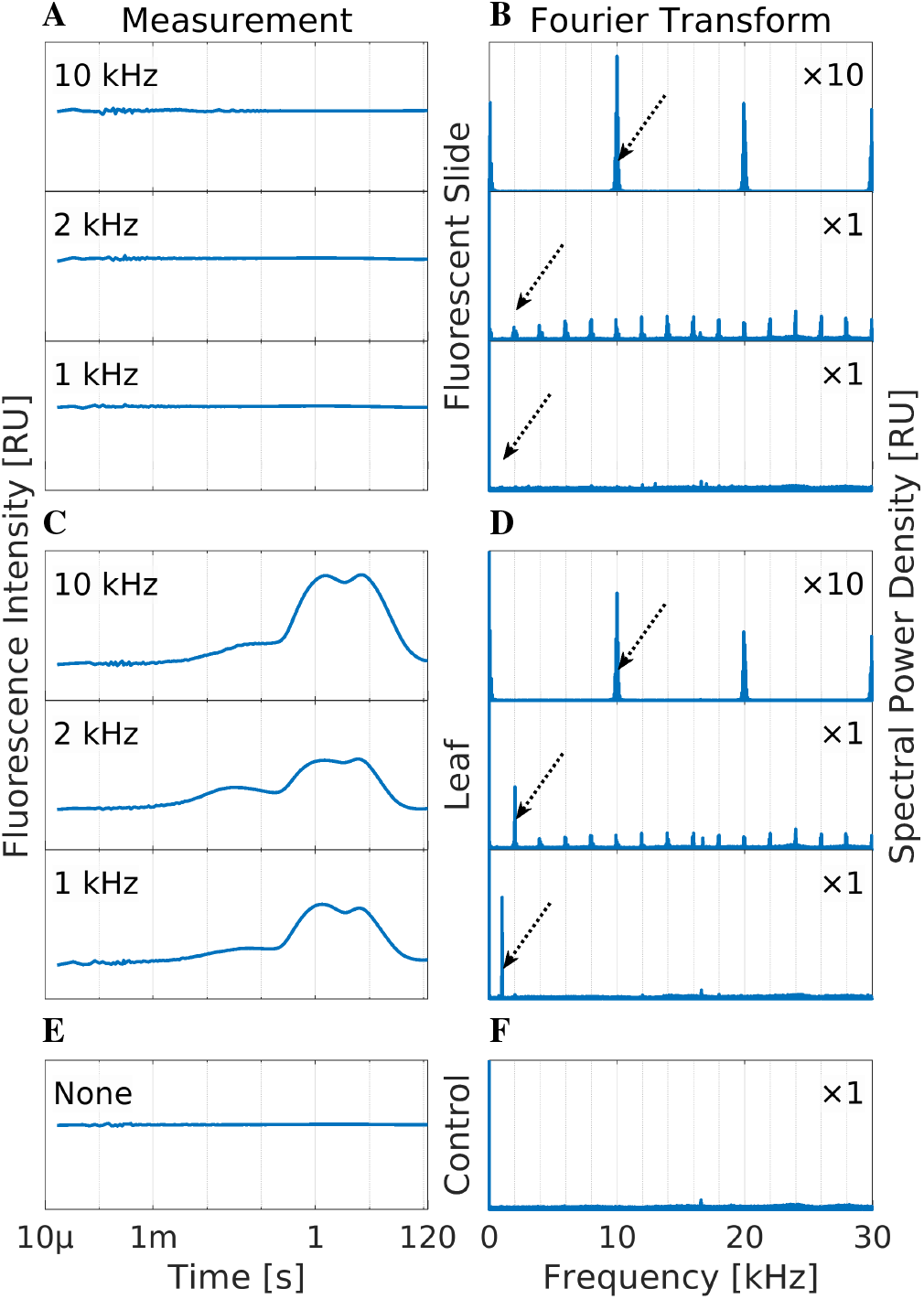
Harmonic light cross-talk at high frequencies. (**A, C, E**) The left column shows the fluorescence intensity traces on a logarithmic time scale. (**B, D, F**) The right column presents their Fourier transforms. (**A**) Fluorescence intensity measurements on the fluorescent slide sample using harmonic light at 10 kHz, 2 kHz, and 1 kHz and (**B**) their Fourier transforms. (**C, D**) The same as above, but measured on the leaf. (**E, F**) Control experiment with the fluorescent slide but no use of harmonic light. The arrows in the right column show the positions of the harmonic light frequencies in the Fourier spectrum. The Y-axis scale of the graphs showing the Fourier spectra for the experiments measured at 10 kHz harmonic light (1_st_ and 4_th_ rows) are ten-times (10) higher than the rest. The remaining graphs in both columns use the same scales.

### C. Harmonic Light Range and Linearity Calibration

The range in which the harmonic LED output was linearly proportional to the setting was determined by a series of measurements described below. An oscilloscope was used to measure the voltage at the output of the photodiode pre-amplifier with the fluorescence slide used as the sample. The harmonic light settings were independently varied as follows: The amplitudes were varied between 8 and 240 and the offsets between 1 and 160. The results are in Fig. 6, where the graphs show the photodiode pre-amplifier output voltages as a function of each setting, the linear regressions of their linear parts, and the residuals. With settings leading to residuals close to zero, the LED output was linearly proportional to the setting value (Table I). This near-linear range spanned from 14 to 180 and 16 to 180 for the amplitude settings of LEDs 1 and 2, respectively. For the offset settings, it ranged from 5 to 100 for both LEDs. At the lower end, the non-linearity stemmed from operating the DACs, operational amplifiers, and DDS circuits near the supply voltage rail. At the top end, the non-linearity was caused by excessive heating of the LED chips, which decreased their light emission efficiency.

**TABLE I.** The settings resulting in the linear operation of the harmonically modulated light sources.

| Parameter | Switch | Minimum | Maximum |
| --- | --- | --- | --- |
| LED 1 Amplitude | A | 14 | 180 |
| LED 2 Amplitude | a | 16 | 180 |
| LED 1 Offset | O | 5 | 100 |
| LED 2 Offset | o | 5 | 100 |

**FIG. 6.**
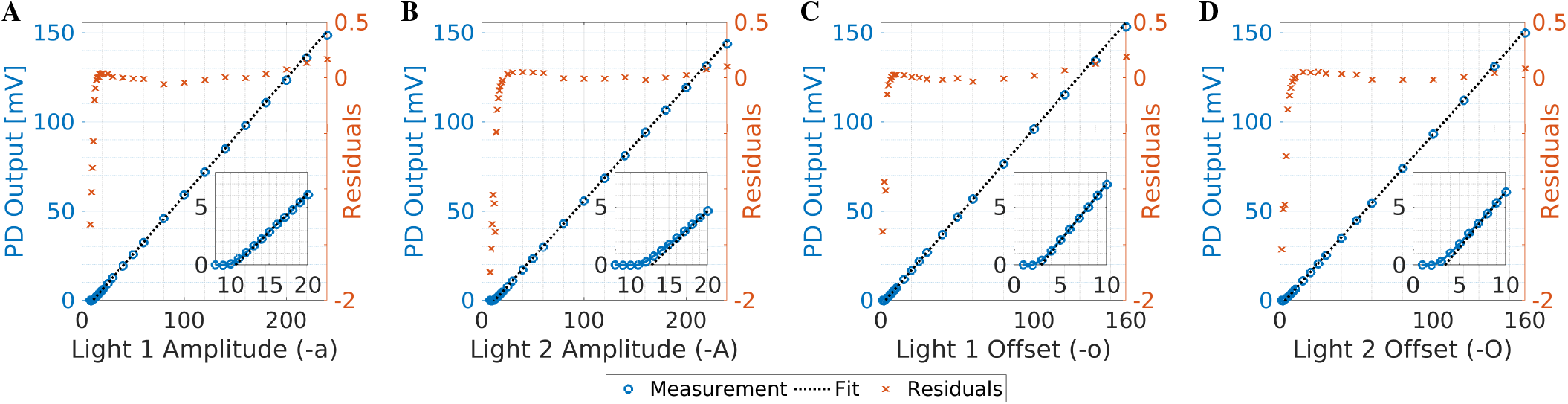
Harmonic light range and linearity. Graphs show photodiode pre-amplifier output voltage dependence on the two harmonic light amplitude and offset settings. Each graph consists of the averaged measured values (blue circles), linear regression fit to the linear part of the data (dotted black line), residuals normalized by 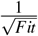 (orange crosses), and an inset zooming onto the non-linear part at lower settings. **A, B** Show graphs of the dependence of the measured voltage on the amplitude settings (-a, and -A, respectively). **C, D** Show graphs of the dependence of the measured voltage on the offset settings (-o, and -O, respectively).

The slopes and zero intercepts of the linear regressions allowed predicting the irradiance at the sample in relative units based on the settings of the offsets and amplitudes of both LED drivers. The equation predicting the photodiode pre-amplifier output voltage (*U*), thus the sample irradiance, was the following:

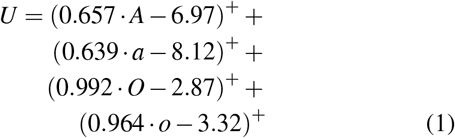

In this Eq. 1 the variables, *A, a, O*, and *o*, were the amplitude of LED 1, amplitude of LED 2, offset of LED 1, and offset of LED 2, respectively. The notation (*v*)^+^ was defined as max{ *v*,0}, meaning that if the value *v* in the parentheses was negative it was replaced by a zero. Notably, the slopes in (Eq. 1) for the amplitude and offset for LED 2 were lower than for LED 1. This suggested that LED 2 produced lower irradiance with the same settings either because of a lower efficiency of the LED chip or imperfect mounting of the collimator lens.

The validity of the prediction of the above equation was tested by 20 different randomly-generated combinations of amplitudes and offsets for both LEDs with linearly increasing expected photodiode pre-amplifier output voltage. The predicted values were compared to the measured values in Fig. 7. The graph shows that the measured values closely matched the predicted values in the range between 20 mV and 300 mV. While the absolute units (mV) of the measurement were dependent on the fluorescent yield of the sample, the important result was that the calibration in Eq. 1 was accurate and the response of the LEDs was linear and behaving as expected in the tested range. In conclusion, the amplitudes and offsets could be combined with the irradiance governed by Eq. 1 and give rise to the expected irradiance.

**FIG. 7.**
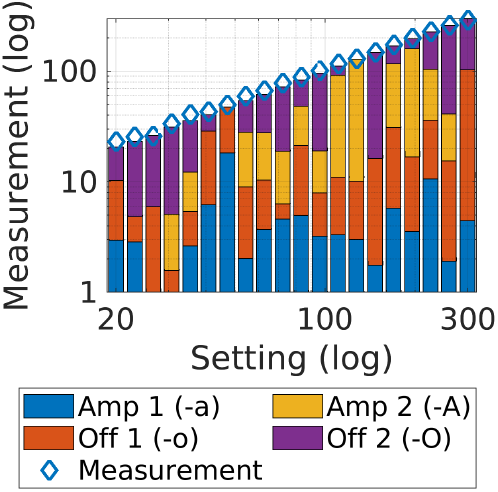
Verification of harmonic light calibration. Combinations of both lights’ amplitude and offset settings were randomly generated to create a linearly increasing light output on a log-log scale. The expected calculated outputs based on these amplitudes and off-sets are determining heights of the bars. The colored subdivisions of the bars are the settings: harmonic light 1 amplitude (Amp 1 (-a), blue), harmonic light 1 offset (Off 1 (-o), orange), harmonic light 2 amplitude (Amp 2 (-A), yellow), and harmonic light 2 offset (Off 2 (- O), purple). Oscilloscope measurements (blue diamonds) were done at the output of the photodiode pre-amplifier during the operation of the Harmonizer with the given settings.

It is important to note that the measurements with given amplitude setting were producing an LED current at the midpoint of the harmonic function. This meant that the LED current at the maximum would be double the amount. Consequently, the maximum setting should not exceed half of the maximum found to be giving a linear LED response, i.e. 90, to ensure linear operation throughout the full cycle of the harmonic modulation. At higher frequencies, where the thermal capacity should keep sufficiently low chip temperature throughout the modulation cycle, the amplitude may be increased further while still retaining a linear operation. Furthermore, the offset and amplitude for each harmonic light are summed to contribute to the resultant LED current. This means the maximum amplitude setting still producing a linear response will be decreased with the applied offset based on the ratio derived from Eq. 1.

### IV. HARMONIC LIGHT MODULATION SHAPE

The harmonic light modulation shape was studied to find the range where it deviates from a harmonic function. The Harmonizer was run with combinations of amplitude (-a) settings ranging from 16 to 240, frequency (-f) settings ranging from 0.1 Hz to 10 kHz, and offset (-o) kept at zero. The measurements were performed using an oscilloscope probe attached to the output of the photodiode pre-amplifier and the fluorescent slide was used as the sample. The results are presented as a set of graphs in Fig. 8 and R-squared values in Table II, which quantify the goodness-of-fit of the harmonic function to the measurement data. The main finding was that at low amplitude settings (*<* 30) the harmonic light intensity waveform deviated from the harmonic shape near its minimum. The residues showed that the amplitude and the shape of the deviation from the harmonic shape remained comparable for in the 0.1 Hz to 1 kHz range for each amplitude setting. At 10 kHz the residuals were worse compared to the lower frequencies. This was not a problem, as according to Fig. 6, frequencies above 1 kHz were best avoided anyway to minimize

**FIG. 8.**
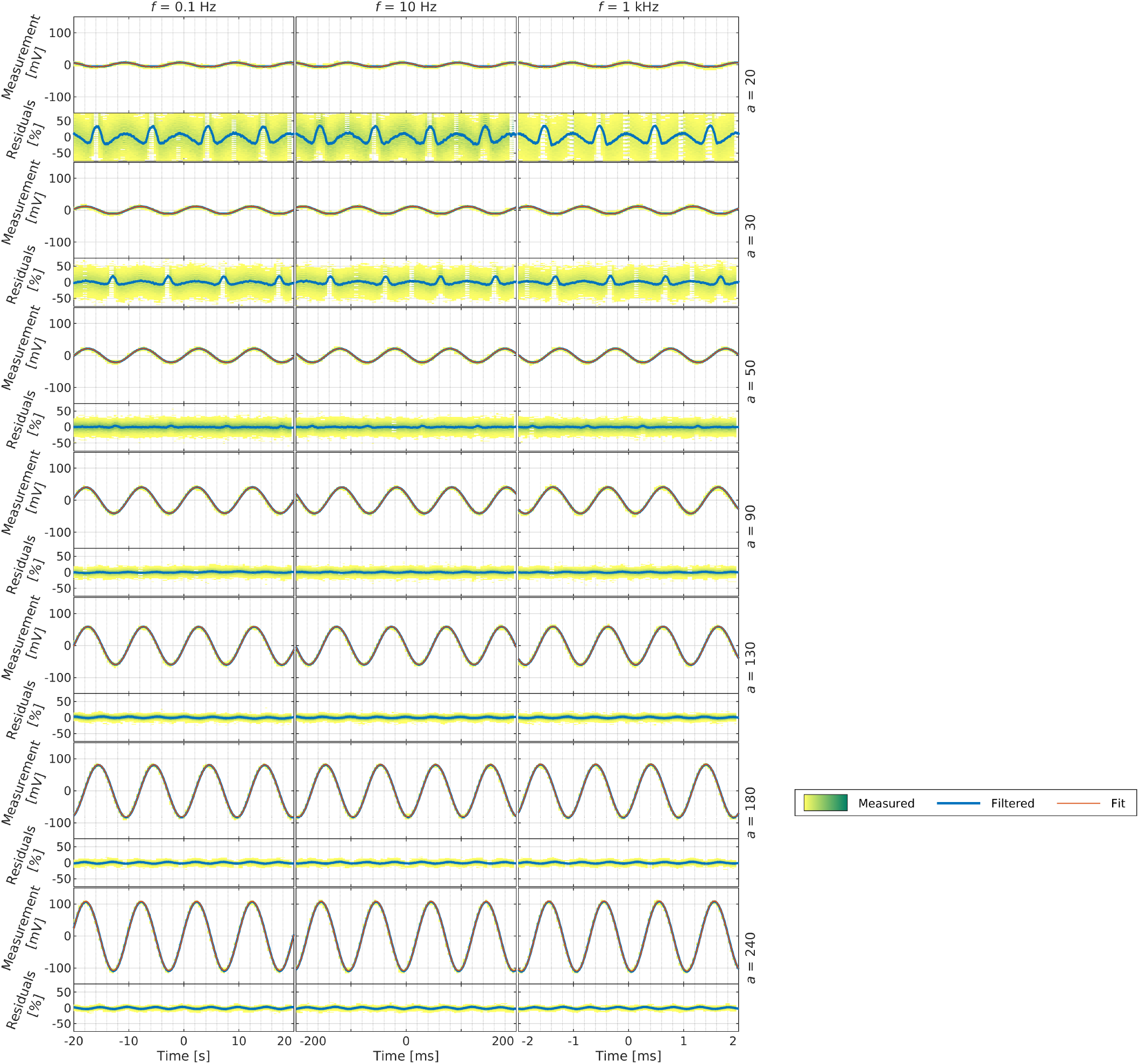
Harmonic shape verification. 21 graph pairs showing the light modulation and residuals as measured by an oscilloscope (measured, yellow-green density map), a moving average (filtered, blue line), and a fitted harmonic function (fit, red line). The top graph in each pair shows the light modulation expressed in millivolts (mV) measured by the oscilloscope. The bottom graph shows the residuals of the fit expressed in percentage of the fitted harmonic function amplitude. The graphs are organized into columns by the modulation frequency (-f) and rows by the modulation amplitude (-a). The frequencies are above the graphs and range from 0.1 Hz to 1 kHz. The amplitudes are to the right of the graphs and range from 20 to 240. Goodness-of-fit of harmonic functions to the measured data are quantified in Table II.

**TABLE II.**
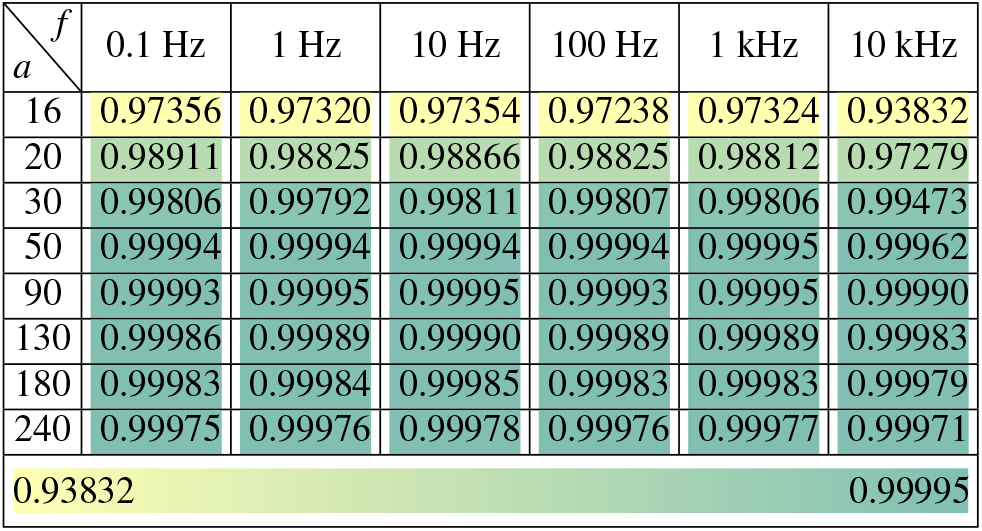
The goodness-of-fit of the harmonic function expressed as R squared. The modulation frequency is in the columns, the amplitude in the rows. The value and the color of each cell denote the R squared value of the fit for the representative measurement with yellow a low R-squared value and green a high R-squared value. The last row contains the lookup table of the cell colors and its range.

harmonic light cross-talk with the measuring light flashes. The color heat map in Table II made the most obvious demonstration of the deviation from the harmonic function shape at the amplitude settings of 16 and 20. Each measurement producing the presented data consisted of many oscilloscope data points (350 000 at 10 kHz, 700 000 at 1 Hz, and 1 400 000 at the remaining frequencies). Since so many datapoints would overwhelm the graphs, they were represented by yellow-green heat maps in Fig. 8. At lower amplitudes, the electronic noise from the Harmonizer, the oscilloscope probe, and the oscilloscope contributed considerably to the final measurements. The harmonic light source was unlikely to contribute measurably to this noise. The noise caused the wider spread of the residue heat maps at the lower amplitude settings. The moving average filtered out this measurement noise and therefore provided a more accurate representation of the actual harmonic light modulation shape compared to the heat maps.

### V. SAMPLE IRRADIANCE CALIBRATION

Up to this point, the Harmonizer was used to characterise its performance. A power meter (PM100A, Thorlabs) equipped with a photodiode (S120VC, Thorlabs) was used to calibrate the sample irradiance by the harmonic LEDs. An iris diaphragm was temporarily fixed to the measurement surface of the Harmonizer. A 3 mm diameter *d* opening was centered on the Harmonizer measurement area. The photodiode was positioned and fixed just above the iris diaphragm and oriented parallel with one of the harmonic light sources to maximize the sensitivity of light detection. Light power *P* incident onto the photodiode was recorded during a Harmonizer measurement with 0.1 Hz frequency and a range of different amplitudes. The measurement was repeated for both harmonic light sources. The power was converted into the units of photosynthetic photon flux density (PPFD) expressed in μmol·m^−2^ s^−1^. For this conversion, the center wavelength *λ* of 460 nm was assumed, the area of the iris opening was scaled by cos(40°) ≈ 0.77 to compensate for the angle *α* between the surfaces of the harmonic LED and the diaphragm

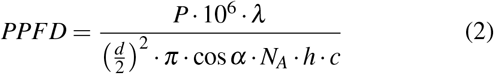

*N*_*A*_ is Avogadro’s constant, *h* Planck’s constant and *c* the speed of light. The results of this calibration are in Fig. 9. The calibration was similar to earlier measurements (Fig. 6) in finding a good linearity, which slightly deviated from the linear response at higher (>130) amplitude settings due to LED chip heating-induced drop in efficiency. The conversion from the amplitude setting to the PPFD irradiance was calculated by fitting a line to the measurements with amplitudes settings in the range between 30 and 130. The resulting conversion from the two amplitude settings (-A and -a) to PPFD values was:

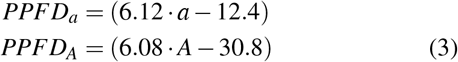

**FIG. 9.**
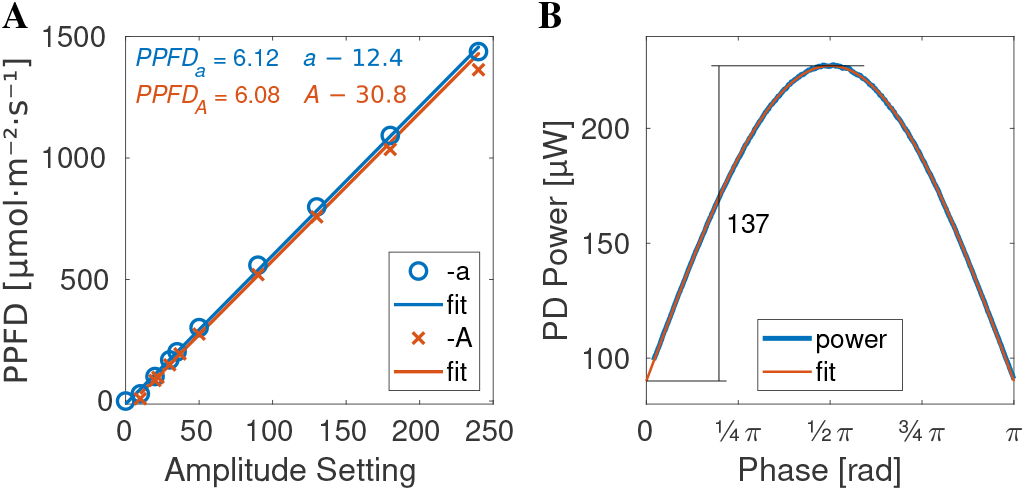
Absolute calibration of harmonic light sample irradiance. **A** Dependence of PPFD amplitude on the amplitude settings -a (blue) and -A (red) with the respective linear regressions (fit). These regressions show that the dependence was linear between ≈ 30 and ≈ 130. The functions describing these linear dependencies are in the top-left corner and Eq. 3. **B** The PPFD amplitude was calculated from the amplitude of a sine function fit to the measured photodiode (PD) incident light power. The amplitudes were converted to PPFD in **A** using Eq. 2.

## VI. PLANT LEAF MEASUREMENT

The Harmonizer was applied to measuring fluorescence of a detached plant leaf with its petiole immersed in water. A series of 12-minutes long measurements were performed in short succession with two different amplitude settings (20 and 35) and 6 frequency settings: 0.0071 Hz (≈ 1*/*141 s), 0.0096 Hz (≈ 1*/*104 s), 0.0110 Hz (≈ 1*/*91 s), 0.0121 Hz (≈ 1*/*83 s), and 0.015 Hz (≈1*/*67 s). The frequencies were selected to cover the range in which nonphotochemical quenching can take place effectively^11^. These amplitudes and frequencies were used either on one or both channels to perform a total of 23 experiments. The data from these experiments were analyzed with the intention to answer the following questions: (i) Does the fluorescent yield curve shape depend on the harmonic excitation light amplitude? (ii) Is a fluorescence yield response to a combination of two harmonically modulated actinic light sources same as the sum of the individual parts? The results presented below demonstrate that the answer is “no” to both questions in the tested scenarios. This means the response of the plant to harmonically modulated actinic light signal is not linear in a mathematical sense and that the time-domain fluorescence induction experiments and frequency-domain experiments yield complementary information about the dynamics of plant photosynthesis.

The first experiment shows how the fluorescence yield changed with harmonic light amplitude. High, medium, and low harmonic light amplitude was used at 0.0110 Hz (≈ 1*/*91 s) frequency and the fluorescence yields were plotted in Fig. 10**A**. The graphs allow making several observations. (i) The plant adapted to the driving frequency over the course of the measurement and this adaptation was slowest and most pronounced with the highest harmonic light amplitude. (ii) The fluorescence yield response was not harmonic. (iii) The peaks of the fluorescence yield were delayed in relation to the peaks in the harmonic light amplitude. These observations are quantified in graphs in Fig. 10**B D**. These graphs characterize five periods of the harmonic light between 200 s and 720 s of the experiments. Fig. 10**B** shows the phase delay between the fluorescence yield peak and the subsequent harmonic light intensity peak (Δφ_*max*_). The phase delay was stable for all three harmonic light amplitudes and the five measured periods. Its values ordered from the highest to the lowest amplitude were (0.591±0.009)*π*, (0.391±0.010)*π*, and (0.292±0.007)*π*. Fig. 10**C** shows the phase delay between the trough of the fluorescence yield and the subsequent harmonic light intensity trough (Δφ_*min*_). The phase delay was dropping for the high harmonic light amplitude during the experiment and seemed to have plateaued around the same value as the phase delay of the troughs for the medium amplitude. The phase delay was stable for the lower two intensities. Its values, ordered from the highest to the lowest amplitude, were (0.65±0.11)*π*, (0.742±0.018)*π*, and (1.761± 0.005)*π*. The amplitudes were increasing during the measurement for all three harmonic light amplitudes and were 72±3, 44.9±0.6, and 27.9±1.1, ordered from the highest to the lowest amplitude. Summarized, the data show that the response of the plant to harmonic light modulation at the 0.0110 Hz frequency differs according to the amplitude. The fluorescence yield response in not harmonic. Between 200 s and 720 s of each experiment, the fluorescence yield response was largely stationary, but at the highest amplitude it was evolving both in the persistently increasing oscillation amplitude and the decreasing phase delay of the signal troughs. The conclusion is that the fluorescence yield response to harmonic light is nonlinear and thus its shape is a function of the harmonic light amplitude.

**FIG. 10.**
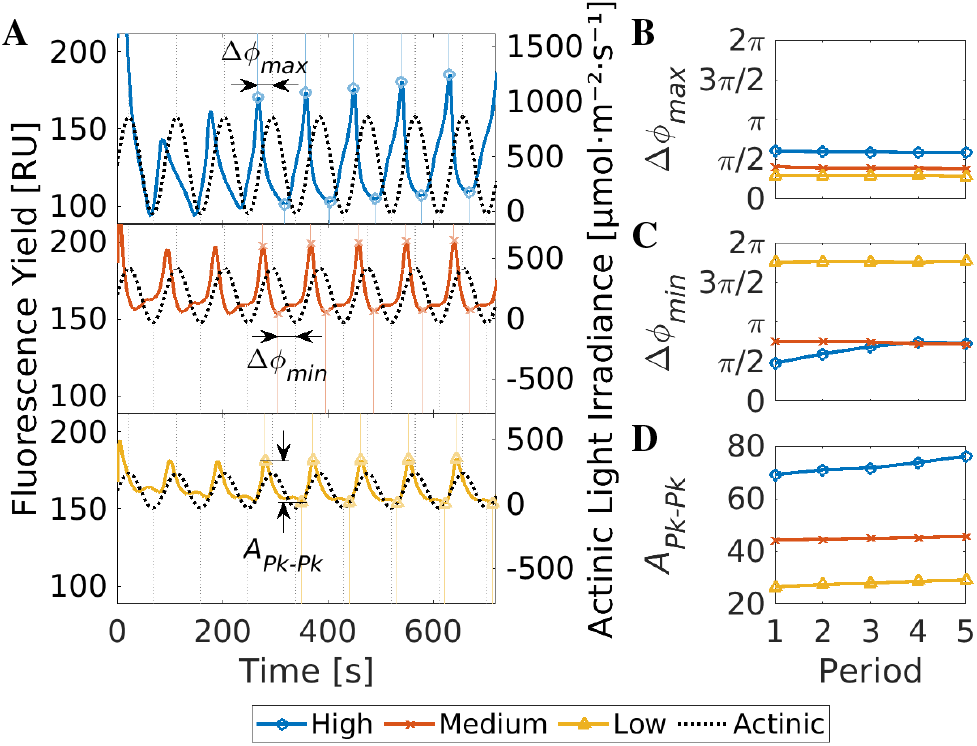
Leaf’s response to different harmonically modulated light intensities. **A** Three harmonic light amplitude settings were used to excite fluorescence with 0.0110 Hz frequency for 720 seconds. The leaf surface irradiances for the settings “High”, “Medium, “Low” were 396, 182, and 91 μmol photons m^*-*2^ s^*-*1^, respectively. The fluorescence yield is plotted in three graphs ordered from high to low harmonic excitation light amplitude. The time positions of the minima and maxima signals are highlighted by vertical dotted lines. The fluorescence yield in the first 200 seconds is ignored to allow for a partial adaptation to the harmonic light modulation. **B** Δ*f*^*max*^ is the phase shift between the fluorescence peak and the subsequent harmonic light maximum as a function of the periods beyond 200 seconds. **C** Δ*f*^*min*^ is the phase shift between the fluorescence yield minimum and the subsequent harmonic light minimum. **D** *A*^*Pk Pk*^ is the amplitude of the fluorescence yield as a function of the periods beyond 200 seconds.

The second experiment tested the question whether the fluorescence yield response to two combined harmonic light sources was same as the sum of its parts. In the five experiments, one harmonic light source frequency was maintained at 0.011 Hz while the other was switched between 0.0071 Hz, 0.0096 Hz, 0.011 Hz, 0.012 Hz, and 0.015 Hz. The amplitudes of the two light sources were set to 20 and 22 to result in equal irradiance. The results are presented in Fig. 11. The results for each combination of harmonic light frequencies are in two graphs forming the rows of the figure. In the left column of each row are the measurements from two harmonically modulated light sources used separately with only one turned on at the time. The time evolution of the two light irradiances is shown by the thin blue (Act fix) and red lines (Act var). The resulting fluorescence yields measured with each of these lights are highlighted by the thick blue (FY fix) and red lines (FY var). The light modulation frequencies are in the top-right corners. In the right column are the two light sources modulated at the same frequencies as in the left column, but applied simultaneously. The thin yellow line (Act comb) shows the time evolution of the combined irradiance from both light sources. The measured fluorescence yield with this combined irradiance is in the thick yellow line (FY comb). A hypothetical sum of the fluorescence yields from the individual use of the light sources, calculated by summing the fluorescence yields shown in the left column, is shown by the thick lavender line (FY fix+var). The graphs show that the response to both harmonic lights at different frequencies simultaneously illuminating the plant leaf leads to a different fluorescence yield compared to the sum of its parts. As with the first experiment, the shape of the fluorescence yield is not harmonic and is different to the harmonic light intensity shape and principal phase shift. In conclusion, the experiments demonstrate that it is impossible to infer fluorescence yield in response to two harmonically-modulated light sources from the response observed with each light source used individually.

**FIG. 11.**
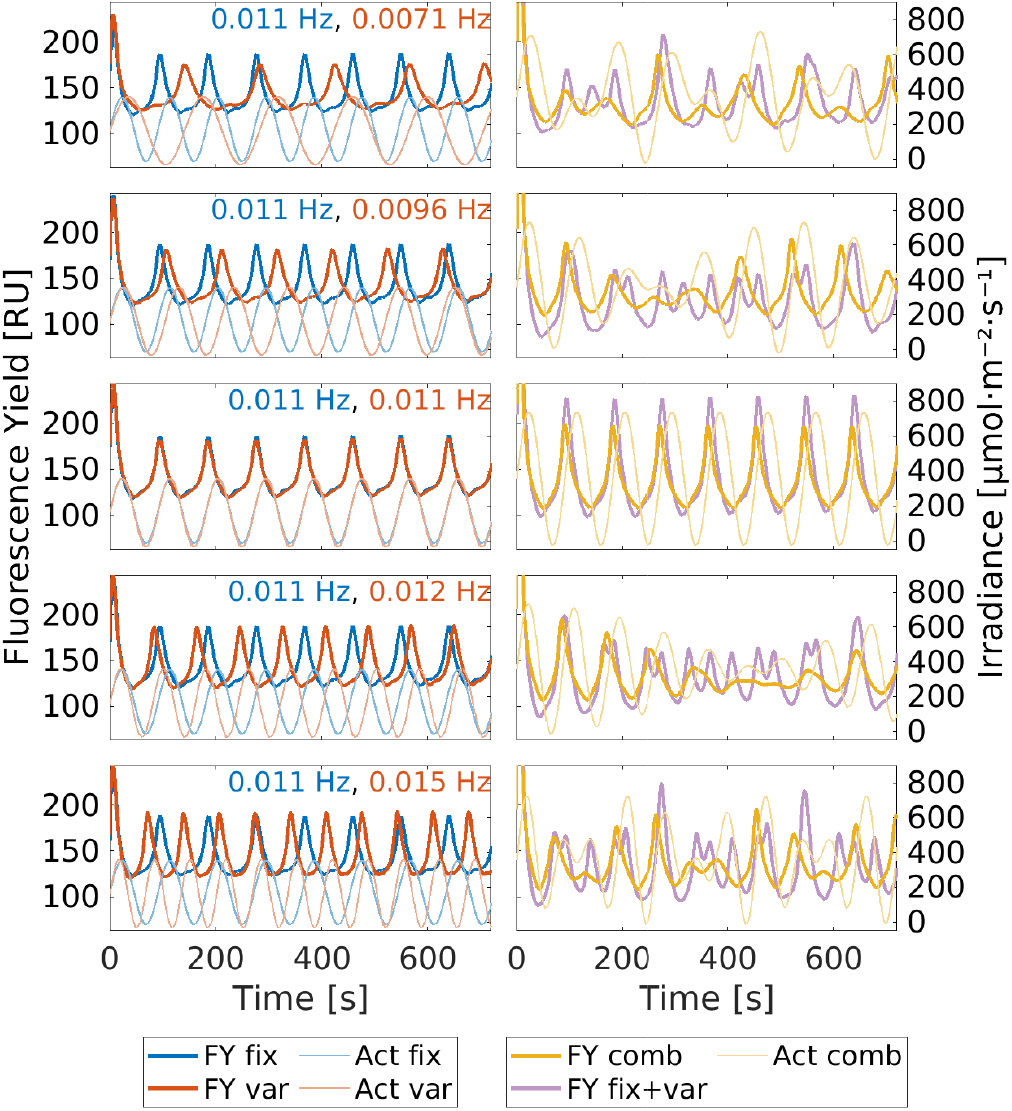
Leaf’s response to two harmonically-modulated light sources. Each row of graphs represents two harmonic light frequencies used in the experiment noted in the top-right corner of the graphs in the left column. One harmonic light frequency is fixed at 0.011 Hz (Act fix) the other is varied (Act var) and the shapes of the harmonic lights’ intensities are in the left graph drawn by the thin lines. The resulting fluorescence yields (FY fix and FY var) for each of these frequencies applied separately are visualized by the thick lines. In the right column is the combined harmonic light irradiance for both light being used simultaneously (Act comb) visualized by the thin yellow line. The measured fluorescence yield (FY comb) for both frequencies applied simultaneously is drawn by the thick yellow line. The hypothetical mathematical sum of the fluorescence yields as a result of the harmonic lights used individually (FY fix+sum) is drawn by the thick purple line.

## VII. DISCUSSION

### A. Instrument and Calibration

The Harmonizer was built from modules connected together to make the instrument. It was designed specifically to perform PAM measurements of plant leaves under harmonic illumination by two independently controlled actinic light sources. It allows measuring changes to the fluorescence yield of plants in response to harmonic light on a background of constant light, a regime particularly suitable to field measurements, where standard PAM requires dark-adaptation. The Harmonizer uses a higher frequency of measuring flashes compared to standard PAM instruments. These measuring flashes causing photochemical reactions, which is undesirable in standard PAM, but here it merely adds to the constant background illumination and vastly improves the achieved signal-to-noise ratio through the averaging of data points in time. The instrument was characterized, operational limits were established and testing on plant leaves was performed. The work has also revealed a number of improvements that could be made to the Harmonizer. The harmonic function shape suffers from distortions at the lower (*<* 30) amplitude setting (Figs. 6, 8). This is due to the way the amplitude is controlled by changing the current at the control input of the sine-wave generator circuit AD9834. A different method for amplitude such as using a digital potentiometer to attenuate the output of the AD9834 is likely to solve this problem. The distortion at higher amplitudes is due to LED chip heating causing drop in its current-to-light conversion efficiency. This problem could be overcome by a employing a feedback circuit that measures the LED light output to compensate for the efficiency changes. This solution could however lead to crosstalk between the two harmonic light sources without a careful design. The Harmonizer harmonic distortion is lowest at PPFD amplitude around 500 μmol photons· m^−2^ s^−1^. This is rather high and could be easily lowered by increasing the feedback resistor in the LED driver modules. This would enable harmonic light operation at lower irradiances. A limitation of the instrument is the measurement duration limited to 12 minutes due to the limited RAM of the Raspberry Pi. This limitation could be overcome by real-time averaging of the measurement data to produce smaller datasets or by employing a Raspberry Pi with more on-board storage RAM.

### B. Plant Measurements

The Harmonizer was built to study a plant’s response to a combination of two harmonic light illuminations. It was used to demonstrate that the response to harmonic excitation is nonlinear. Using the same frequency and changing the amplitude of the harmonic light caused differently shaped fluorescence yield responses at the settings used. It was observed that the plant adaptation to the harmonic light is gradual and slower with higher (amplitude 400 μmol photons· m^−2^ s^−1^) irradiance. Combining two harmonic lights of different frequencies did not lead to the same fluorescence yield response as the sum of the individual parts. This finding is of principal importance because it means that in the range of tested experimental conditions, the time-domain and frequency-domain measurements are not mutually related by Fourier transform and represent complementary information on plant photosynthesis. The typical time-domain experiment identifies fluorescence induction^13^ during dark-to-light transition whereas a Bode plot in the frequency-domain characterizes photosynthetic dynamics in the light-acclimated plant.

The Harmonizer demonstrated the capability to perform high signal-to-noise measurements not possible using PAM instruments currently available on the market, offering a detailed design that can be easily reproduced at a cost that is a fraction of commercial instruments and further modified for further frequency-domain studies. The presented experiments merely demonstrated the possibilities, but are by no means exhaustive. Measurements on a steady background illumination could be performed to mimic measurements in field conditions. Response to harmonic light could be studied at much wider range of frequencies (1 kHz−0.0001 Hz). The response to varying phase shift between the two excitation frequencies could be studied. The two harmonic lights could have different wavelengths. A red harmonic light could be used alongside or instead of blue excitation or a near-infrared and blue harmonic lights could be combined to study the effect of photosystem I sinking the electron pool created by photosystem II.

These studies are of imminent importance to understanding the performance of plants in dynamically changing light and improving yield potential of our major crops^14^.

## ACKNOWLEDGMENTS

Funding to King’s College was provided by the Engineering and Physical Sciences Research Council (UK), grant number EP/X525571/1.

## DATA AVAILABILITY STATEMENT

The data, source code, design files for 3D printing of the Harmonizer optomechanical parts, and electronics schematics are available from the King’s College London research data repository, KORDS, at https://doi.org/10.18742/c.7180416^15^. The original printed circuit board (PCB) designs will be made available with an accompanying disclaimer on request addressed to the authors. The PCB design files were not added to the supplementary materials because post-production modifications were made to them. These changes were marked in the schematics, but updated PCBs were never designed and produced

Appendix A: Source Code

The source code is provided in the repository^16^. The archive file Harmonizer.code.zip contains a folder arduino, which contains the source code main.ino for the Arduino UNO. Arduino must be electrically disconnected from the Harmonizer before programming. Arduino UNO was programmed using the desktop IDE version 1.8.19. A second folder within the archive is called rpi, which contains the Python scripts for the Raspberry Pi, to control the Harmonizer. The only script required by the user of the Harmonizer is rpi/main/main.py. The archive also contains the file Installing_RPi_to_support_SPI_and_I2C.pdf. This PDF explains how to configure the RPi to operate with the Harmonizer.

The rest of this appendix describes how to use the main.py script to control the Harmonizer. It can be run alone, using the default value, or with any of the switches described in Table III. Next is an example command to run the Harmonizer. The explanation follows below.

~~~
python3 main.py -a 35 -A 0 -m 20 --name LEAF
~~~

**TABLE III.** main.py script to run the Harmonizer. Switches to the script can change the experimental parameters and they are described in this table. All switches are optional and if they are not called, the default value (third column) will be assumed. The switch can either take the short form (first column) or the long form (second column). The script will ensure the supplied values are within the permitted ranges. The long form distinguishes the two light sources as blue and red. This is a legacy issue, as both harmonic lights in the Harmonizer are blue (460 nm). The experiment always completes with saving a file, the prefix to the file can be chosen using the -n or --name switch.

| Short switch | Long switch | Default | Range | Description |
| --- | --- | --- | --- | --- |
| -n | --help |  |  | Show help message without running an experiment |
| -a | --blueAmplitude | 40 | 0 – 255 | First harmonic light amplitude |
| -o | --blueOffset | 0 | 0 – 255 | First harmonic light offset |
| -p | --bluePhase | 0 | –180 – 180 | First harmonic light phase |
| -f | --blueFrequency | 1 | 0.001 – 10000 | First harmonic light frequency |
| -A | --redAmplitude | 40 | 0 – 255 | Second harmonic light amplitude |
| -O | --redOffset | 0 | 0 – 255 | Second harmonic light offset |
| -P | --redPhase | 0 | –180 – 180 | Second harmonic light phase |
| -F | --redFrequency | 1 | 0.001 – 10000 | Second harmonic light frequency |
| -m | --pulseAmplitude | 10 | 0 – 255 | Measuring flash amplitude |
| -s | --seconds | 10 | 2 – 10000 | Experiment duration in seconds |
| -n | --name | test | a-Z0-9_-. | Name of experiment |
|  | --graph, --no-graph | --graph |  | Produce or not produce a graph |

`python3 main.py` ran the script. The switches that followed set the experimental parameters. `-a 35` set the amplitude of the first harmonic light to 35. `-A 0` set the amplitude of the second harmonic light to 0, which means the light remained turned off during the experiment. `-m 20` set the amplitude of the measurement flashes to 20. `--name LEAF` was an example of a long form switch and the value `LEAF` was going to become part of the produced file name. The remaining parameters that were not set stayed at the default values summarized in Table III.

When this script finished, it produced a file and a graph (Fig. 2**A**, inside the screen). The file-name contained all important metadata of the experiment. The filename structure is described below: `2022_11_09-23_37_35---LEAF a35-o0-p0_0.0-f1. 0_1.0133-A0-O0-P0_0.0-F1.0_1.0133-m20-s10.bin`. The first numbers were the date and the time of the experiment completion in YYYY_MM_DD-HH_MM_SS format. This was followed by the name of the experiment `LEAF`, which was specified by the `--name` switch. Then were the amplitude `-a35` and offset `-o0` of the first harmonic light.

This was followed by `-p0_0.0`. The `-p0` meant that the phase of the first harmonic light was set to zero (0). The number behind the underscore (_) was the actual value of the phase (0.0). Since the phase and frequency cannot be set to any arbitrary value, the value behind the underscore was the actual experimental value, closest possible to the desired one. The first light frequency `-f` was set to 1.0 Hz, but actually was 1.0133 Hz, the value behind the underscore. Next came the amplitude (`-A0`), offset (`-O0`), and frequency (`-F1_1.0133`) of the second light, followed by the amplitude of the measuring flashes (`-m20`), and the experimental duration (`-s10`).

